# Evaluation of combined root exudate and rhizosphere microbiota sampling approaches to elucidate plant-soil-microbe interactions

**DOI:** 10.1101/2025.10.23.683011

**Authors:** C Escudero-Martinez, E Y Browne, H Schwalm, M Santangeli, M Brown, L Brown, D M Roberts, A.M. Duff, J Morris, PE Hedley, P Thorpe, J Abbott, F P Brennan, D Bulgarelli, T S George, E Oburger

## Abstract

- Deciphering the root exudate-driven interplay between plants and the rhizosphere microbiota is essential for understanding plant adaptation to the environment and future-proofing crop production. However, sampling root exudates and rhizosphere soil remains challenging due to the low throughput and destructive nature of the process.
- We used the staple crop barley [*Hordeum vulgare*] as a model to benchmark different sampling approaches for simultaneous exudation and microbiota profiling of soil-grown plants.
- Exudate profiles and total dissolved organic carbon exudation rates were consistent across different sampling approaches, whereas root biomass, root morphology measurements, and organic nitrogen exudation varied.
- High-throughput amplicon sequencing and quantitative PCR (qPCR) of phylogenetic markers and nitrogen cycle-selected genes revealed a protocol-specific footprint in the composition and abundance of rhizosphere bacterial and fungal microbiota. Yet, on average, 75% of microbes enriched in, and differentiating between, barley rhizosphere and unplanted soil controls were recovered across all the sampling approaches evaluated.
- Our results demonstrated that, under the tested conditions, different sampling approaches produced comparable microbiota and exudation patterns, enabling the integrated study of root exudation and microbial profiles from the same plant. The observed differences across sampling approaches must be considered according to the experimental scope.

## Introduction

Plant-soil-microbe interactions in the rhizosphere, or the interface between roots and soil, play a pivotal role in plant health and growth performance. Therefore, the rhizosphere can be regarded as an ‘extended’ root phenotype (de la Fuente Cantó *et al*., 2020). Soluble root exudates, i.e., plant metabolites released via the root into the surrounding soil, are key drivers of biogeochemical processes in the rhizosphere as they shape the abundance, function, and composition of the microbial communities populating the rhizosphere, designated ‘rhizosphere microbiota’ (Anderson *et al*., 2024). Harnessing beneficial rhizosphere traits, including root exudation and the rhizosphere microbiota, is a critical step toward more sustainable agriculture (Schneider & Lynch, 2020; Brooker *et al*., 2022; Oburger *et al*., 2022; Colombi *et al*., 2024). Significant progress has been made in phenotyping (i.e., measuring the plant growth, development, and physiology arising from interactions between genotypes and environment) of root architectural traits, paving the way for including these traits in breeding programs (Paez-Garcia *et al*., 2015). However, investigating complex rhizosphere traits such as plant-soil-microbe interactions remains challenging, especially at a throughput relevant to plant breeding.

Diverse toolboxes to study root exudation (Oburger & Jones, 2018), the rhizosphere microbiota, (Alegria Terrazas *et al*., 2016) and additional rhizosphere traits such as the rhizosheath formation, pH and elemental gradients have already been established (George *et al*., 2008; Neumann *et al*., 2009; Oburger & Schmidt, 2016). However, most techniques are often tailored for in-depth studies and/or handling a limited number of genotypes and/or treatments, preventing a full appreciation of how plant genetics impacts rhizosphere traits. Likewise, most techniques and sampling approaches focus on individual aspects of the rhizosphere traits, e.g., either exudates or the microbiota, preventing a holistic insight into feedback processes. For instance, root exudation and microbiota composition, and their subsequent impact on plant performance, are often inferred from independent samples rather than the same plant specimen due to handling complexity (Jin *et al*., 2024; Zhalnina *et al*., 2018).

Phenotyping complex rhizosphere traits, such as root exudation, is methodologically challenging and labour-intensive. Efforts to quantify root exudation from soil-grown plants are confounded because collecting rhizosphere soil solution does not enable accurate determination of root exudation rates, as the samples also contain dissolved native soil (Oburger & Jones, 2018). Furthermore, exuded metabolites can immediately interact with the soil matrix (sorption) and be instantly decomposed by the rhizosphere microbiota (Oburger *et al*., 2009; Oburger *et al*., 2011; Oburger *et al*., 2016). To circumvent soil matrix interactions during the sampling process, root exudates are often collected from plants grown in (sterile) hydroponics or inert, artificial growth substrates (e.g., Kawasaki *et al*., 2018; Lopez-Guerrero *et al*., 2022; Aleksza *et al*., 2024; Otxandorena-Ieregi *et al*., 2024). Artificial growth conditions also enable better control of microbial activity, especially during the exudate sampling process. Microbial decomposition in non-sterile exudate sampling approaches has been repeatedly shown to alter metabolite concentrations (Valentinuzzi *et al*., 2015; Otxandorena-Ieregi *et al*., 2024). Therefore, controlling microbial activity during the sampling process is critical if the aim is to obtain unbiased information about the quantity and diversity of plant-exuded metabolites (Oburger & Jones, 2018). However, the ecological relevance of artificial growth environments, which are distant from field conditions, may fail to recapitulate rhizosphere processes (Oburger & Jones, 2018). If the aim is to investigate plant-soil-microbe interactions, working with soil-grown plants is essential. Among protocols using soil-grown plants, the soil-hydroponic-hybrid technique, which combines plant soil growth with a short hydroponic exudate collection period (Oburger *et al*., 2014; Santangeli *et al*., 2024), is probably the most applicable approach for investigating root exudation in the context of rhizosphere processes at an ecologically relevant scale. This protocol requires (i) the excavation of an intact root system, followed by (ii) careful soil removal by root washing (to avoid root damage), (iii) a short root damage capture/osmotic adjustment period, and (iv) 2 – 4 hours (h) of hydroponic exudation sampling. The replicate throughput of this protocol is sufficient for medium-scale experiments (∼100 samples), including genotypes and/or treatments. Also, despite the sudden change in environment from soil to solution, the root cell environment will still reflect natural soil growth conditions, especially when the sampling period is short.

Like root exudates, experimental protocols have been developed to sample the rhizosphere soil for microbiota analysis, designated ‘rhizosphere fractionation.’ Typically, the bulk soil assumed to be unaffected by root activity is removed by gently ‘shaking’ the roots until only the soil tightly adhering to the root remains. This root-adhered soil is then defined as the rhizosphere soil (Escudero-Martinez *et al*., 2022; Oyserman *et al*., 2022; Chang *et al*., 2025). In a subsequent step, the rhizosphere soil is harvested from the roots. This can be achieved by gently brushing roots (e.g., with a sterile toothbrush) (Lucas *et al*., 2018; Fukuda *et al*., 2022). However, this rhizosphere sampling approach is time-consuming, may cause root damage, and carry brushed root cellular material into the sample. To address these limitations, we previously developed a two-step sampling approach that separates soil from roots more effectively. Firstly, after the bulk soil is removed, rhizosphere-coated roots are submerged in a buffer solution, followed by vortexing to facilitate the rhizosphere separation. Finally, the rhizosphere soil is centrifuged to create a rhizosphere soil pellet for microbial DNA extraction. This high-throughput method, capable of handling ∼100 samples, has been successfully applied to studies focused on the rhizosphere microbiota, as it consistently retrieves a set of microbes significantly different from the unplanted soil (Maver *et al*., 2021; Chang *et al*., 2025) (Terrazas *et al*., 2022; Reid *et al*., 2024).

Both rhizosphere microbiota and root exudate analyses have already been successfully combined with other root and shoot traits, including root morphology, shoot nutrient content, and root gene expression (Escudero-Martinez *et al*., 2022; Malacrinò & Bennett, 2024; Santangeli *et al*., 2024). However, despite the mechanistic link between root exudates and rhizosphere microbiota, these traits have typically been sampled from different plant specimens within the same study (e.g. Zhalnina *et al*., 2018; Escudero-Martinez *et al*., 2022) due to trait-specific sampling requirements. In practice, this means that root exudates are collected from living, intact plants, while rhizosphere soil is usually sampled after destructive excavation of a part of the root system. Integrating the sampling of root exudates and rhizosphere microbiota from the same plant specimen will (i) avoid the uncertainty of linking different rhizosphere traits collected from two individual plants (even within the same experiment), (ii) increase the number of treatments that can be investigated within one experiment as only half the number of plants are needed to sample both exudates and rhizosphere microbiota and (iii) deliver a fundamental understanding on the root exudate metabolites-microbiota feedback processes.

Hence, the objective of this study was to develop, assess, and critically evaluate different sampling approaches that enable the collection of root exudates and the rhizosphere microbiota from the same soil-grown plant. As a proof of concept, we used the staple crop barley [*Hordeum vulgare*], one of the reference plants for microbiota investigations in crops (Alegria Terrazas *et al*., 2020; Jiang *et al*., 2025). By adapting two well-established sampling approaches, the soil-hydroponic hybrid exudate sampling approach (Oburger *et al*., 2014; Otxandorena-Ieregi *et al*., 2024) and the root shaking/vortexing rhizosphere microbiota sampling approach (Robertson-Albertyn *et al*., 2017; Alegria Terrazas *et al*., 2020; Escudero-Martinez *et al*., 2022; Terrazas *et al*., 2022), we tested different procedures of (i) bulk soil removal (rinse vs. shake), (ii) the rhizosphere harvest from roots (dipping vs. vortexing) and the effect of the final rhizosphere fraction collection (gravitational sedimentation vs. centrifuging), followed by a 3 h hydroponic exudate collection period (Fig. 1). We compared the impact of the different approaches on plant biomass, root morphology, root exudate quantity (total C exudation rates) and the metabolic profile of root exudates (non-targeted metabolomic fingerprinting using Reversed-Phase Liquid Chromatography–Mass Spectrometry (RPLC-MS), as well as on rhizosphere microbiota composition by metabarcoding of bacteria/archaea (16S rRNA gene), and fungi (ITS), absolute quantification of these microbial communities and the abundance of functional microbial nutrient cycling genes (quantitative PCR).

**Fig. 1:**
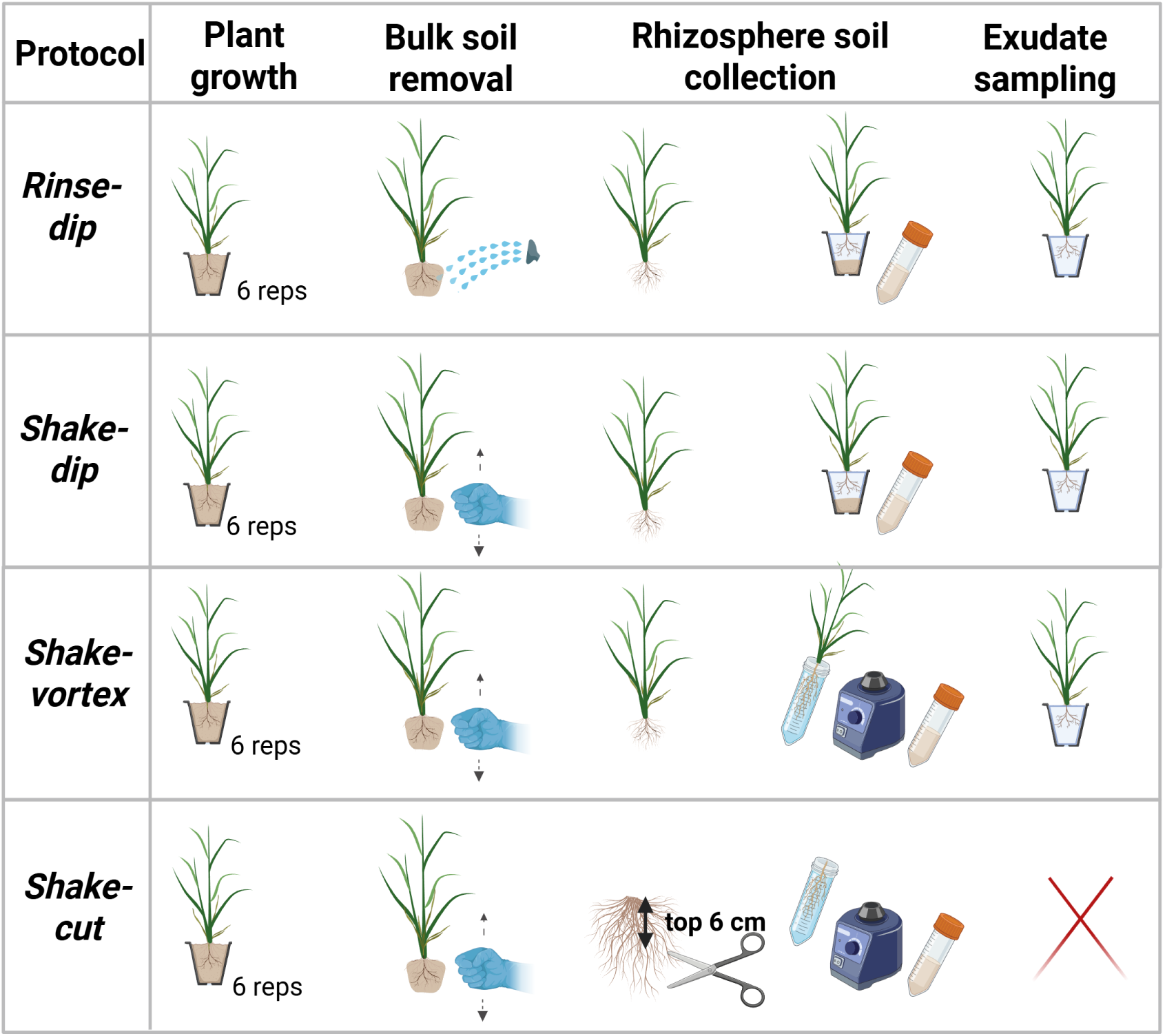
Schematic representation of the steps involved in the different rhizosphere sampling protocols *Rinse-dip*, *Shake-dip*, *Shake-vortex*, and *Shake-cut*.

## Materials and Methods

### Plant growth

Barley (*Hordeum vulgare*) plants from the genotype cv. Barke were used to evaluate the different sampling approaches (Fig. 1). Barley seeds were surface sterilised by washing them in 70% ethanol (30 s) followed by 5% sodium hypochlorite (15 min). Sterilised seeds were rinsed four times with autoclaved deionised water and pre-germinated on Petri dishes containing a semi-solid 0.5% agar solution. After 2 days, germinated seeds with similar rootlet size were sown in individual 400 mL pots filled with 300 g of a sieved (10 mm) reference agricultural soil previously used for barley-microbiota investigations, designated ‘Quarryfield’ (Invergowrie, Scotland, UK, 56° 27′ 5” N 3° 4′ 29” W; Sandy Silt Loam, pH 6.2; Organic Matter 5%;) (Robertson-Albertyn *et al*., 2017; Alegria Terrazas *et al*., 2020; Maver *et al*., 2020; Escudero-Martinez *et al*., 2022; Terrazas *et al*., 2022). Planted pots were arranged in a randomized design *n* = 6 replicates per sampling approach, plus *n* = 6 unplanted (bulk) controls for *Rinse-dip* and *Shake-dip*, and an additional n = 6 unplanted controls for *Shake-vortex* and *Shake-cut*. Watering was performed daily to 80% field capacity using sterilised deionised water by weighing the pots. Two weeks after planting, plants were supplied weekly with a modified Hoagland’s solution where mineral nitrogen was reduced to 25% (Terrazas *et al*., 2022). Plants were grown until stem elongation, Zadok’s stage 30-35; ∼5-week post-transplant) in a glasshouse at the James Hutton Institute, UK under the following controlled environmental conditions: 18/14 °C (day/night) temperature regime with 16 h daylight that was supplemented with artificial lighting to maintain a minimum light intensity of 200 µmol quanta m^−2^ s^−1^.

### Testing different sampling approaches for combined rhizosphere soil collection and exudation sampling

Considering the scope of this methodological study, we report the tested experimental approaches in a protocol-like manner to facilitate the applicability and reproducibility of the different sampling approaches. An overview of the four different sampling approaches is presented in Fig. 1.

#### 1. Protocol Rhizosphere *Rinse-dip*

##### 1.1. Bulk soil removal

The rooted soil blocks were carefully removed from the pots, ensuring the plant remained intact. They were then rinsed under a stream of tap water (i.e., wet bulk soil removal via rinsing) until only a thin rhizosphere soil layer attached to the roots remained.

##### 1.2. Rhizosphere soil collection - part I

Once the rhizosphere soil layer was visible, the root system was repeatedly dipped into a deionised water (DI) container to dislodge the rhizosphere soil gently. The entire root system was submerged in this solution. The dipping process was repeated using a second container with fresh DI water to maximise rhizosphere soil collection. This soil slurry represented the rhizosphere fraction. Both dipping solutions were pooled into one container, and the rhizosphere fractions were left for sedimentation for at least 2 h at 4 °C. The dipping solution volume was adapted to the target plant’s species root system size. In this barley experiment, we used a 2 x 250 mL dipping solution.

##### 1.3. Exudate sampling

In the meantime, roots were subjected to one final, gentle rinse and placed into a fresh container with DI water, awaiting further processing. Before submerging the roots into the final exudate sampling solution, they were placed into a fresh container with the same composition as the exudate sampling solution. This aimed to capture metabolites released from damaged cells and promote a root osmotic adjustment. After 5 min, the roots were removed and gently placed on tissue paper for a few seconds to capture water droplets. This exudate sampling solution was discarded. The roots were then transferred to the final container with 80 mL of exudate sampling solution. The exudate sampling solution contains autoclaved DI containing 5 mg L^-1^ Micropur (Roth, Katadyn), as a broad-spectrum bactericide to avoid microbial decomposition of exuded metabolites during the exudate collection (Otxandorena-Ieregi *et al*., 2024). The sampling solution containers were wrapped with aluminium foil to protect the roots from light and placed back into the greenhouse for 3 h (Aulakh *et al*., 2001). We aimed for the root (dry weight) to sampling solution volume (L) ratio (RSVR) to be in the range of 2-4 to avoid any set-up driven biases (Otxandorena-Ieregi *et al*., 2024).

##### 1.4. Rhizosphere soil collection - part II

While the plants were in the greenhouse for exudate collection, containers with the rhizosphere soil slurry were taken from the 4 °C cold room. After gravitational sedimentation for at least 2h at 4 °C, the supernatant was discarded by pouring until a 50 mL sample was obtained. Six supernatant samples were kept as ‘supernatant controls’ for the *Rinse-dip* and *Shake-dip* protocols. The concentrated 50 mL samples were centrifuged at 1500 g for 20 min at room temperature, and the remaining supernatant was removed. The resultant rhizosphere pellet was flash-frozen in liquid nitrogen and stored at −70 °C.

##### 1.5. Plant tissue harvest & processing of exudate samples

After an exudate collection period of 3 h, plants were removed from the containers. The containers with the exudate sample solution were closed and stored at 4 °C until further processing. Then the plant material was harvested. First, roots and shoots were separated. The shoot fraction was collected in paper envelopes for dry weight determination by drying at 70 °C for 2 days. The root system was briefly blotted with paper before fresh weight determination. Then, the root system was divided into two aliquots. This was achieved by cutting the root system into four equal transversal sections (S1-S4) along the longitudinal axis. Each of the four sections was further evenly divided into two halves by manually separating the roots (SX-right and SX-left). These sections were then combined into two aliquots following the longitudinal root axis to ensure the representativeness of the entire root system (S1-S4 right and S1-S4 left): one section was dedicated to dry weight determination (2 days at 70 °C) and the second section for the analysis of root morphological parameters by WinRHIZO (See ‘Root morphology analyses’) and was stored in 70% ethanol until processed.

Immediately after plant tissue harvest, exudate samples were filter sterilised (0.2 μm acetate filters, Chromafil®) and aliquoted as follows: 30 mL for total dissolved organic carbon (DOC) analysis, 30 mL for non-targeted metabolite analysis, and the remaining ∼20 mL as backup. Samples were frozen at –20 °C, freeze-dried in an Alpha 1-2 LD plus lyophiliser (Sci-Quip), and then shipped to BOKU, Vienna, Austria, for root exudate analysis.

#### 2. Protocol Rhizosphere *Shake-dip*

##### 2.1 Bulk soil removal

The rooted soil blocks were carefully removed from the pots, ensuring the plant remained intact. Roots were gently shaken to remove loosely bound soil particles until only the rhizosphere soil tightly adhered to the roots remained, i.e., dry bulk soil removal by shaking. This soil was considered the rhizosphere fraction.

##### 2.2. Rhizosphere soil collection - part I

The intact root system with the adhering rhizosphere soil was repeatedly dipped in DI water, following the same procedure from this point on as described in *1.2. Protocol Rhizosphere Rinse-dip*.

##### 2.3. Exudate sampling

Same procedure as described in *1.2. Protocol Rhizosphere Rinse-dip*.

##### 2.4. Rhizosphere soil collection - part II

Same procedure as described in *1.2. Protocol Rhizosphere Rinse-dip*.

##### 2.5. Plant tissue harvest & processing of exudate samples

Same procedure as described in *1.2. Protocol Rhizosphere Rinse-dip*.

#### 3. Protocol Rhizosphere *Shake-vortex*

##### 3.1. Bulk soil removal

Dry bulk soil removal by shaking – same procedure as described in *2.1. Protocol Rhizosphere Shake-dip*.

##### 3.2. Rhizosphere soil collection

the above ground plant material was separated from the root system. The intact root system, along with its adhering rhizosphere, was placed into a sterile 50 mL Falcon tube containing 20 mL of phosphate-buffered saline (PBS). Samples were then vortexed for 30 s, and the rhizosphere soil was sedimented for 2–3 min, incubated in ice. The roots were transferred to a new 50 mL Falcon tube with 20 mL PBS, in which the samples were vortexed again for 30 s to separate the remaining rhizosphere soil from the roots. The two falcon tubes containing the rhizosphere fraction were combined into a single tube, with the combined sample representing the rhizosphere fraction. This sample was then centrifuged at 1500 g for 20 min at room temperature. After centrifugation, the supernatant was discarded, and the pellet was flash-frozen in liquid nitrogen and stored at −70 °C.

##### 3.3. Exudate sampling

Same procedure as described in *1.1. Protocol Rhizosphere Rinse-dip*.

##### 3.4. Plant tissue harvest & processing of exudate samples

Same procedure as described in *1.1. Protocol Rhizosphere Rinse-dip*.

#### 4. Protocol Rhizosphere *Shake-cut*

This protocol represents a well-established reference for 16S rRNA gene and ITS metabarcoding sequencing that has been frequently used in previous investigations. (Maver *et al*., 2021; Escudero-Martinez *et al*., 2022; Terrazas *et al*., 2022) However, sampling a section of the root system (the root system uppermost 6 cm) represents a destructive approach that cannot be combined with root exudation sampling.

##### 4.1. Bulk soil removal

Dry bulk soil removal by shaking – same procedure as described in *2.1. Protocol Rhizosphere Shake-dip*.

##### 4.2. Rhizosphere soil collection - part I

The root system was sectioned to retain just the uppermost 6 cm of the seminal root system. Then, the samples were vortexed and centrifuged as described in *3.2. Protocol Rhizosphere Shake-vortex*.

##### 4.3 Exudate sampling

No exudates were collected as the root system did not remain intact.

##### 4.4. Plant the harvest & processing of exudate samples

In this protocol, only the shoots were harvested and processed as described in *1.1. Protocol Rhizosphere Rinse-dip*.

### Unplanted soil collection

#### Bulk soil collection corresponding to protocols Rinse-dip and Shake-dip

In same the area explored by the roots in the planted plots, the same soil volume obtained from the planted samples was collected from the unplanted pots. The soil was collected with a spatula and mixed with 250 mL of DI water. The bulk soil suspension was processed as in *Rinse-dip and Shake-dip: Rhizosphere soil collection - part II*.

#### Bulk soil collection corresponding to protocols Shake-vortex and Shake-cut

In the same area explored by the roots, the same soil volume obtained from the planted samples was collected from the unplanted pots in the planted plots. The soil was collected with a spatula and mixed with 30 mL of PBS. The bulk soil suspension was further processed as in the *Shake-vortex* and *Shake-cut Rhizosphere soil collection*.

### Root morphology analysis

The root sections stored in 70% ethanol were spread out in a clean, scratch-free plastic tray filled with water. Roots were arranged to avoid overlapping and any meniscus bubbles. Images were scanned in black and white at 600 dpi on an Epson 12000 XL flatbed scanner (Watford Hets England WD17 1JA).

Scanned images were analysed using WinRHIZO Pro™ software (WinRHIZO Pro 19, Regent Instruments Inc., Canada) to obtain root morphological traits. Root diameter categories were set at increments of 0.5 mm. The detection threshold was set at “automatic.”

Several root morphology measurements were recorded (Table S1). For calculations, we first normalised the WinRHIZO outputs by the total root dry weight. The specific root length (SRL) was computed using the equation root length /root dried weight. Normality was checked using the Shapiro–Wilk test, and one-way ANOVA and post hoc Tukey’s HSD were used to assess differences in root morphology.

### Microbiota analysis

Total DNA was extracted from the rhizosphere and unplanted soil samples for the four different sampling approaches using FastDNA™ SPIN kit for soil (MP Biomedicals, Solon, USA) following the manufacturer’s instructions eluted at 35 ul with deionised and distilled water and subject to targeted amplification of microbiota phylogenetic markers.

### Amplicon sequencing library preparation: 16S rRNA gene

16S rRNA from rhizosphere and unplanted soil preparations were generated using the 515F-806R primer pair (Caporaso *et al*., 2012) (Supplementary Information 1). These PCR primer sequences were fused with Illumina flow cell adaptor sequences at their 5′ termini. The 806R primer contains a 12-mer unique ‘barcode’ sequence used for each sample, enabling multiplexed sequencing of several samples in a single pool.

For each bulk and rhizosphere sample, a total of 50 ng of DNA was subjected to PCR amplification using the Kapa HiFi HotStart PCR kit (Kapa Biosystems, Wilmington, USA). The individual PCR reactions were performed in a 20 µL final volume and contained: 4 µL of 5X Kapa HiFi Buffer, 10 µg Bovine Serum Albumin (BSA) (Roche, Mannheim, Germany), 0.6 µL of a 10 mM Kapa dNTPs solution, 0.6 µL of 10 µM solutions of the individual PCR primers, 0.25 µL of Kapa HiFi polymerase. Reactions were performed using the programme found in Supplementary Information 1. For each primer combination, a no-template control (NTC) was included. Individual PCR reactions were performed in triplicate, and 2 independent sets of triplicate reactions per barcode were included to minimise PCR bias.

Before purification, 6 µL aliquots of individual replicates and the corresponding NTCs were inspected on 1.5 % agarose gel to detect amplification and/or any contamination. Samples with the expected amplicon size and no detectable contamination in NTCs on the gel were used for successive library preparation. Individual replicates of the PCR amplicons were then pooled per replicate and purified using the Agencourt AMPure XP Kit (Beckman Coulter, Brea, USA) with 0.7 times the volume of the sample beads. Purified samples were quantified using a Qubit Fluorometer with the dsDNA HS assay (Invitrogen, Thermo Fisher Scientific, USA). Once quantified, individual barcoded samples were pooled into a new single tube in an equimolar ratio to generate the library.

Paired-end Illumina sequencing (2 × 150 bp reads) (Maver *et al*., 2021) was performed using the Illumina MiSeq system. Library pool quality was assessed using a Bioanalyzer (High Sensitivity DNA Chip; Agilent Technologies) and quantified using a Qubit and qPCR (Kapa Biosystems, Wilmington, USA). Amplicon libraries were spiked with 15 % of a 4 pM phiX control solution. The resulting high-quality libraries were run at 10 pM final concentration.

### Amplicon sequencing library preparation: ITS region

For amplicon sequencing a 2-step PCR method was used PCR 1 involved the amplifying the ITS2 region of the rhizosphere and bulk soil preparations using the 86F and 4R primer pairs (Supplementary Information 1), each reaction was carried out in a 25 μL volume containing 12.5 μL of KAPA HiFi HotStart ReadyMix Taq (Roche Sequencing), 0.2 μM forward and reverse primers, and 2.5 μL of the template at 5 ng/uL. Cycling conditions for PCR1 are found in Supplementary Information 1. PCR1 was performed with three technical replicates per sample, which were then pooled for purification with 0.6 uL Ampure XP beads (Beckmann Coulter, UK) per uL of DNA followed by elution in 15 μL TE buffer. Next, barcodes were added to the amplicons using the unique dual index (UDI; Integrated DNA Technologies, Germany) (PCR2). Amendment of UDI indexing primers was performed in a 50 μL reaction volume containing 25 μL of KAPA HiFi HotStart ReadyMix Taq, 5 μL of PCR 1 template, and 5 μL of UDI. PCR conditions are found in Supplementary Information 1. Amplicons were purified using the Ampure XP beads as for PCR1. Product concentrations were measured using the Qubit fluorometer dsDNA HS assay (Thermofisher, Ireland). The products were then pooled in equimolar concentration, the library quality was checked using the Bioanalyzer (High Sensitivity DNA Chip; Agilent Technologies), and concentrations were quantified using the Qubit fluorometer. Paired end reads (2 x 300bp) were sequenced on the Illumina NextSeq 2000 system using the P2 flow cell (Teagasc sequencing facility, Ireland).

### Amplicon sequencing reads processing

Reads quality assessment DADA2 version 1.22 (Callahan *et al*., 2016) and R 4.1.0 (R-Core-Team, 2021) were used to generate the ASVs following the ‘DADA2 Pipeline Tutorial (1.10)’ and taxonomic identification against the SILVA 138 database for 16S rRNA and the UNITE v2023 database for ITS (Quast *et al*., 2013). The creation of the Phyloseq object is explained in detail in Maver *et al*., 2021. Subsequently, sequences classified as ‘Chloroplast’ or ‘Mitochondria’ from the host plant were pruned *in silico*. Further filtering criteria were applied for the 16S rRNA gene library; samples with less than 10,000 reads and low-count ASVs were pruned from the Phyloseq object (at least 20 reads in 2% of the samples). The 16S rRNA and ITS datasets were rarefied at equal sequencing depth across samples (64,000 reads for the 16S rRNA gene and 200,000 reads for the ITS). The resulting 16S rRNA gene Phyloseq object was used for downstream analyses, containing 2,376 individual ASVs and 39 samples. The ITS Phyloseq object contained 2,139 individual taxa and 19 samples.

### Calculation of alpha-, beta-diversity indices, and differential abundance

For consistency, the same pipeline was applied to both 16S rRNA and ITS sequences. Alpha-diversity richness was estimated as described in (Maver *et al*., 2021). Beta-diversity analysis was carried out by calculating the dissimilarities among microbial communities using the rarefied data with the Bray-Curtis index as described in (Maver *et al*., 2021). *DESeq2* (Love *et al*., 2014) was used to perform microbial differential abundance analysis to identify genera differentially enriched between pairwise comparisons by Wald test (False Discovery Rate, FDR < 0.05). Microbial taxa significantly enriched in rhizosphere specimens compared to their cognate unplanted soil were used as a proxy for the plant’s effect when comparing sampling approaches and were visualised with the package *UpSetR* (Conway *et al*., 2017).

### qPCR

Before molecular analysis, potential inhibitors in the samples were tested as per Duff *et al*., 2022 (Supplementary Information 1). Briefly, if the plasmid showed reduced amplification compared to the spiked control, the sample was considered to have inhibitors. None of the samples in this study showed evidence of inhibitors and further cleaning of samples was not undertaken.

Following inhibition testing and sample clean up, qPCR was used to quantify the absolute abundance of bacterial 16S rRNA gene and fungal ITS2 copies, as well as nitrification and denitrification genes: *AmoA* and *nosZ* clade II, respectively. A ten-fold serial dilution of the standard was prepared, covering seven concentration points ranging from 10^8^ to 10^2^ gene copies per ng. qPCR was performed in a 10 μL reaction volume containing 5 μL Takyon™ Low ROX SYBR 2X Master Mix blue dTTP (Eurogentec, Ireland), 2 μL of DNA template (1 ng/ μL), and 1.5/0.2 μM each of the primers. Primer concentration, sequences, and cycling conditions are found in Supplementary Information 1. Five samples per sampling approach were analysed with qPCR, and each sample was run with three technical replicates. An NTC was also run for each plate. A positive control of known quantity was also included for the 16S rRNA, ITS and nosZ clade II to confirm accuracy of the assay, a positive control was not used for the AOA qPCR assay due to lack of control availability.

The resulting average gene copies per ng DNA of the three technical replicates for each sample were assessed for normality using a Shapiro-Wilk test. One-way analysis of variance (ANOVA) was performed on each gene, followed by a post hoc Tukey test using *car* and *agricolae* packages in R 4.3.2 (R Core Team, 2023). The residuals of each ANOVA model were visually assessed for homoscedasticity and normality with a residual and a quantile-quantile test. Statistical outliers, i.e., those points that statistically affect the output of each model, were identified using a Cook’s distance test.

### Root exudate analysis and data evaluation

Total dissolved organic carbon (DOC) concentrations and total nitrogen bound (TNb) concentrations were determined using a liquid Total Organic Carbon (TOC) elemental analyser (Elementar VarioTOC, Germany). For the analysis, the freeze-dried aliquot (original sample volume 30 mL) was resuspended in 15 mL of type 1 laboratory-grade water. To determine total dissolved organic N (DON), inorganic N concentrations (only NO_3_ ^-^, no NH ^+^ was detected) were analysed using the spectrophotometric assay from (Hood-Nowotny *et al*., 2010) and NO_3_ ^-^ concentrations were subtracted from TNb. Total dissolved organic carbon (hereafter C) and total dissolved organic nitrogen (hereafter N) exudation rates (nmol cm^-2^ RSA h^-1^) were calculating by dividing the absolute quantity of C and N (nmol) in each sample by the total root surface area (cm²) of the respective barley plant and by the sampling period (3h).

For non-targeted metabolomic analysis, the second freeze-dried exudate sample aliquot (originally 5 mL) was re-suspended in 1.0, 1.5, or 2.0 mL of LC-MS grade water, depending on the original sample DOC concentration, and aiming for a final DOC concentration of around 85 mg C L^-1^. Re-suspensions were filtered (4 mm; 0.2 µm cellulose acetate, Nalgene™, USA) and non-targeted metabolomic analysis was conducted using a 6230b Time-of-Flight mass spectrometer (TOF-MS) with a dual Jetstream ESI interface (Agilent Technologies, Santa Clara, USA) coupled with an Infinity II UHPLC system (Agilent Technologies, Santa Clara, USA). The separation was performed by Reversed-Phase Liquid Chromatography (RPLC), using an Atlantis T3 C18 column (2.1 x 150 mm, 3 µm particle size, Waters Corporation, Milford, USA) as described in Lohse *et al*. (2023). Briefly, chromatographic separation was carried out using gradient elution at a flow rate of 200 μL min^−1^. Mobile phase A consisted of LC-MS grade water with 0.1% v/v formic acid, and mobile phase B was 100% methanol. The initial gradient condition of 100% A was maintained for 2 minutes, then decreased to 60% A over the next 8 minutes and held constant for an additional 2 minutes. This was followed by a cleaning step with 100% B for 2.1 minutes, a rapid decrease to 0% B, and a re-equilibration period of 5.9 minutes at 100% A. The injection volume was 5 μL, and the column temperature was maintained at 40 °C. Mass spectra in the range of 90 to 1700 m/z were recorded in negative (−) polarity, utilising the 2 GHz extended dynamic range mode with a spectral acquisition rate of 2 Hz. Data were acquired with the MassHunter acquisition software (version 10.1, Agilent Technologies, Santa Clara, CA), while MassHunter Profinder B.10.00 software (Agilent Technologies, Sta Clara, CA) was used for peak picking and chromatographic deconvolution with a Batch Recursive Feature Extraction (BRE, small molecules/peptides) workflow (for details see also SI of Lohse *et al*. (2023). Samples were injected in a randomised sequence and quality control samples (QCs, prepared by combining equal volumes of each sample) were injected every 5 samples to monitor the stability of the TOF-MS system. The first four injected samples were excluded by further data evaluation due to pressure instability of the binary pump during the measurement. Features showing a relative standard deviation higher than 25% in the QCs, and features with an intensity of less than ten times the blank average, were excluded from further analysis. Thereafter, raw intensities were scaled with a sample-specific factor calculated as pre-concentration fold x root surface area, followed by unit variance scaling. Finally, multivariate statistical analyses such as PCA and PERMANOVA were performed to investigate the sampling approach’s impact on the metabolomic data.

## Results

### Different sampling approaches have an impact on root biomass and morphology, but not root exudate quantity or metabolic profile

We evaluated the impact of the different sampling approaches, summarised in Fig. 1, on plant biomass (roots and shoots) and root morphology. We used these parameters as indicators of plant integrity. For the *Shake-cut* sampling approach, total root weight, morphology, or root exudate parameters were not measured as roots were partially destroyed during sampling.

As expected, the shoot biomass did not differ in the protocols evaluated (ANOVA; P-value = 0.0684). Conversely, we observed a significant decrease in the recovered root biomass in the following order: *Rinse-dip* ≥ *Shake-dip* ≥ *Shake-vortex*, but only *Rinse-dip* and *Shake-vortex* were significantly different (ANOVA; P-value= 0.005, Fig. 2a, b). Root morphology parameters such as total root surface area (RSA, cm²), average root diameter (mm), and total root length (cm) followed the same trend as the root biomass (Table S1). However, the specific root length (SRL) was comparable across the sampling approaches.

**Fig. 2:**
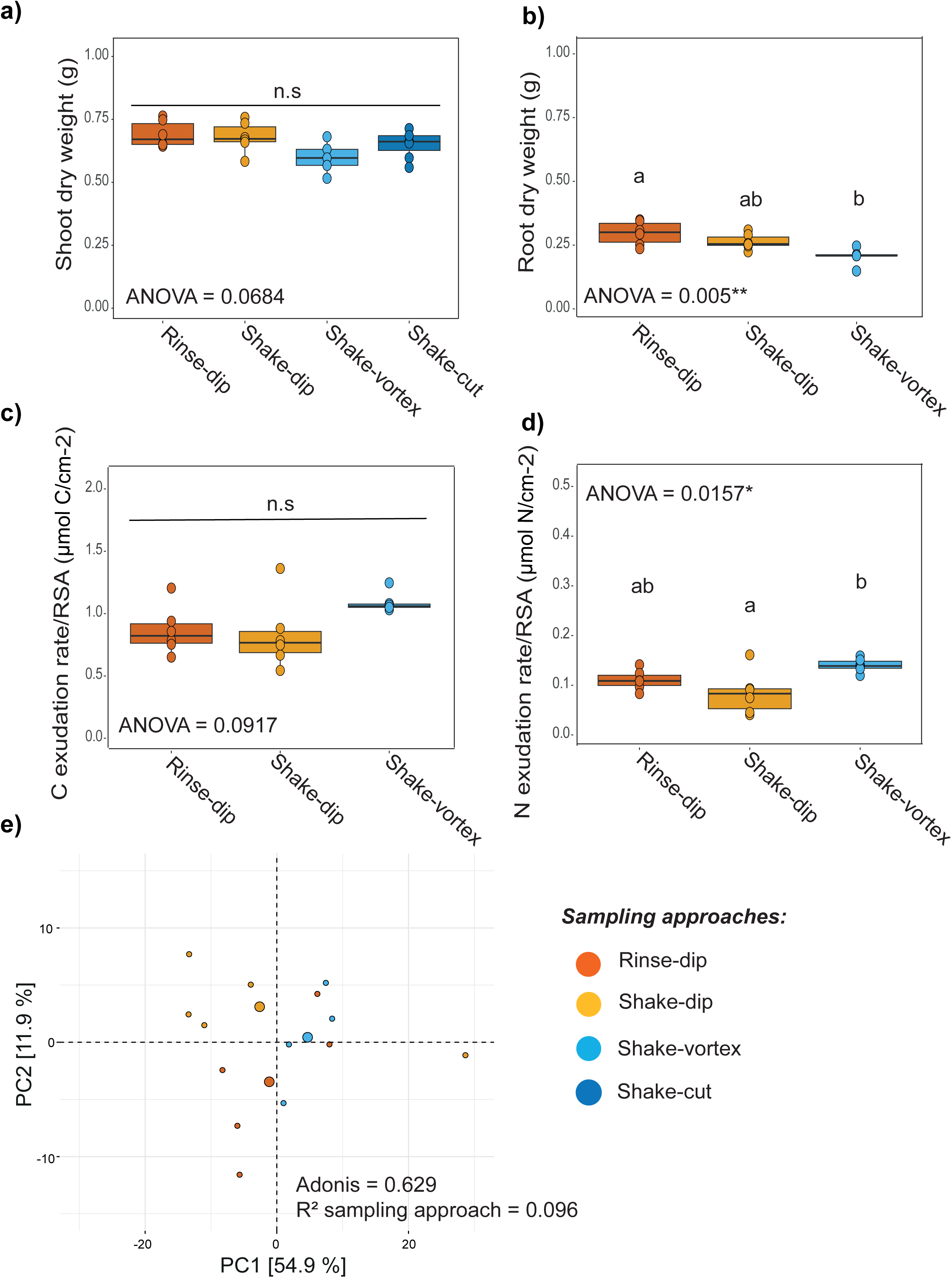
Biomass and exudation across sampling approaches: Boxplot depicting the dry weight biomass of shoot (a), roots (b), Carbon exudation rate (c), and Nitrogen exudation rate (d). In each panel, individual dots depict individual biological replicates. Upper and lower edges of the box plots represent the upper and lower quartiles, respectively. The bold line within the box denotes the median. Letters indicate significant differences following an ANOVA and post-hoc Tukey test (p-value < 0.05). d) Principal component analysis (PCA) of exuded metabolite features detected via non-targeted metabolomics analysis. In the PCA plot, large dots represent group centroids, while small dots indicate individual samples of the four sampling approaches depicted in distinct colours. Adonis was used to test for significance between the sampling approaches (p-value > 0.05). R^2^ indicates the proportion of variance explained by the sampling approach. *Rinse-dip* (orange), *Shake-dip* (yellow), *Shake-vortex* (light blue), and *Shake-cut* (blue).

An important factor for harvesting root exudates is a constant root (dry) weight-to-sampling solution volume (L) ratio (RSVR) (Otxandorena-Ieregi *et al*., 2024). Due to the lower root biomass in *Shake-vortex* and a constant exudate sampling volume, we observed a significantly (p =0.0023) lower RSVR in *Shake-vortex* (2.57 ± 0.16 g root dwt L-1, mean±SE) in comparison to *Rinse-dip* (3.71 ± 0.24) and *Shake-dip* (3.29 ± 0.16) (Fig. S1).

C root exudation rates did not differ between the different sampling approaches, with an average 0.93±0.05 µmol C cm^-2^ RSA h^-1^ (mean±SE) (ANOVA; P-value = 0.0917, Fig. 2c). However, N root exudation rates were significantly affected (ANOVA; P-value = 0.0157, Fig. 2d). *Rinse-dip* N exudation rates (0.11±0.01 µmol N cm^-2^ RSA h^-1^, mean±SE) were comparable to *Shake-dip* and *Shake-vortex*, yet *Shake-dip* (0.08±0.02) N exudation rates were significantly lower than *Shake-vortex* (0.14±0.01).

Metabolomic fingerprinting on the root exudates revealed no significant differences among sampling approaches (Adonis test, P-value = 0.629), further supported by the lack of protocol-wise sample organisation in the ordination (Fig. 2e). Nevertheless, we observed that *Shake-dip* and *Shake-vortex* exudate composition showed more consistent exudate profiles, as indicated by the tighter clustering within those groups (Fig. 2e).

### All sampling approaches distinguish rhizosphere microbial communities from those of unplanted soil

Absolute quantification of bacterial (16S rRNA gene) rhizosphere communities did not identify significant differences between sampling approaches (ANOVA P-value > 0.05, Fig. 3a). Conversely, a significant sampling approach effect was observed on the abundance of the ITS phylogenetic marker (ANOVA P-value = 0.0219, Fig. 3 b). *Shake-vortex* displayed greater ITS copy numbers compared with *Shake-dip*. *Shake-vortex* approach was also associated with greater variability in microbial gene abundances regardless of the taxonomic marker quantified (Fig. 3a, b, Table S2).

**Fig. 3:**
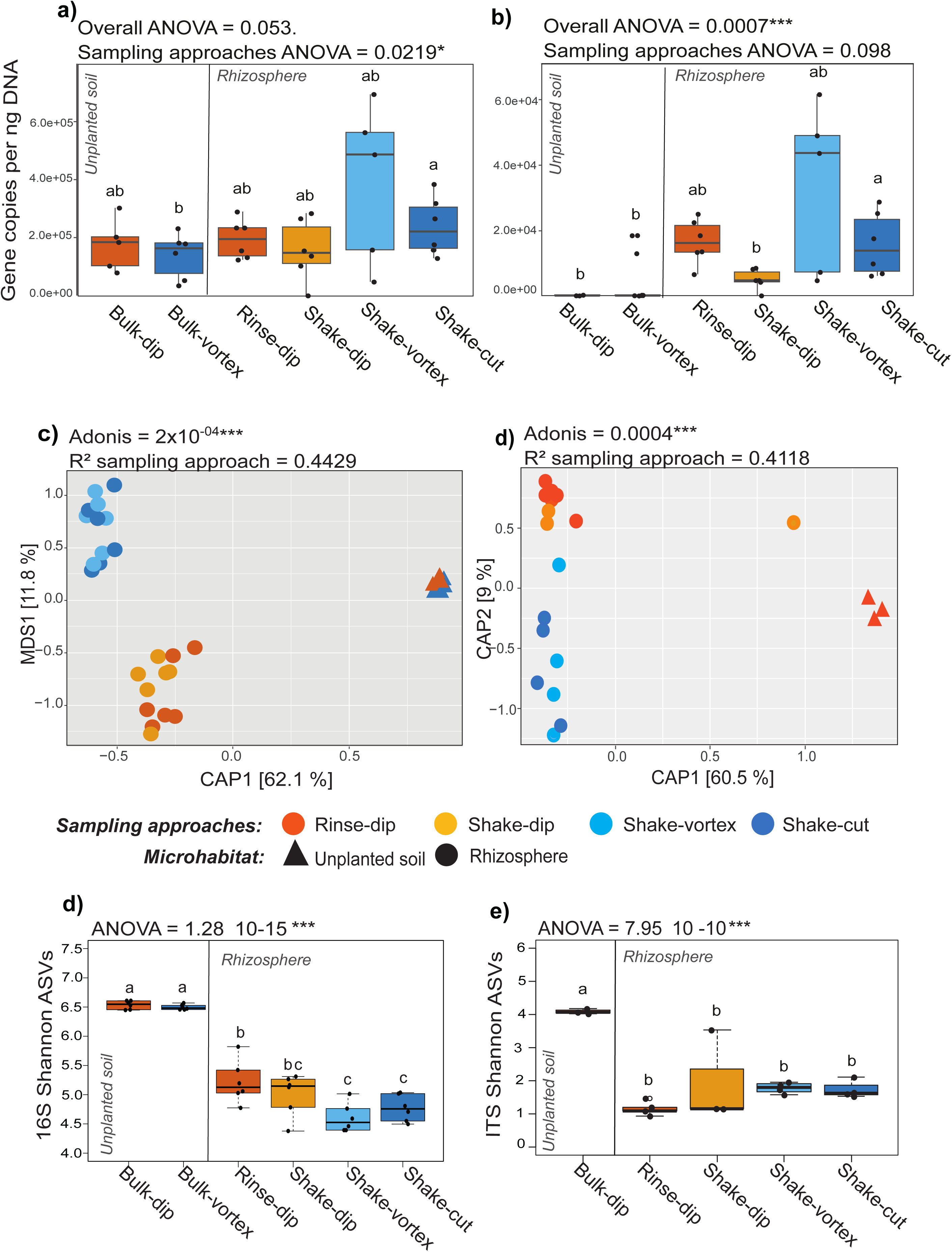
Microbial quantification and diversity: (a) 16S rRNA gene (bacteria) and (b) ITS (fungi) gene copies per ng of DNA in unplanted soil and rhizospheres obtained with different sampling approaches. Box plots depict the total copy numbers of each taxonomic marker per ng of DNA (16 rRNA gene or ITS). Individual dots depict individual biological replicates. Different letters denote significantly distinct groups (ANOVA and post hoc Tukey HSD test Canonical Analysis of Principal Coordinates (CAP) computed on Bray-Curtis dissimilarity matrix for the different rhizosphere sampling approaches and the unplanted bulk soil controls for (c) bacteria 16S rRNA gene and d) fungal ITS data. Sample type is depicted by colours and shapes. Individual dots in the plot denote individual biological replicates of the different sampling methods or the unplanted soil samples. The number in the plot depicts the proportion of variance (R2) explained by the factor ‘Sampling approach’ on rhizosphere samples. The asterisks associated with the R2 value denote its significance in the Adonis test (p-value < 0.05). Note that the y-axis measures Metric Multidimensional Scaling (MDS) and the x-axis represents Canonical Analysis of Principal Coordinates (CAP) ordination, differing one order of magnitude in (c). Boxplots representing the alpha-diversity index Shannon in (e) bacteria 16S rRNA gene and (f) fungal ITS data for the sampling approaches and the unplanted bulk soil controls. In each panel, individual dots depict individual biological replicates. Empty dots are outliers. Upper and lower edges of the box plots represent the upper and lower quartiles, respectively. The bold line within the box denotes the median. Letters indicate significant differences following ANOVA and post-hoc Tukey test (p-value < 0.05). In boxplots, orange bulk soil samples are the controls for *Rinse-dip* and *Shake-dip*, while blue bulk soil samples are the controls for *Shake-vortex* and *Shake-cut*.

Beta-diversity, or between-sample microbial diversity, confirmed a significant rhizosphere effect for bacteria/archaea and fungi, irrespective of the sampling approach. Rhizosphere communities across all the sampling approaches were significantly different from the unplanted soil as shown by sample separation along the x-axis of the Canonical Analysis of Principal coordinates (CAP) Adonis,16S rRNA R2= 78%, P-value = 2×10^-4^, ITS R2=86 %, P-value = 4×10^-4^, 5,000 permutations, Fig. 3c and d). Note that hereafter for fungi we used as unplanted reference the bulk soil of the ‘dipping’ approaches as the ‘vortexing’ bulk soil did not pass the ITS sequencing quality filters. Beta diversity within the rhizosphere sampling approaches showed differences in bacterial composition across all the approaches, except for the comparison between *Shake-vortex* and *Shake-cut*, with no significant variation (Adonis test, Table S3a). Sampling approaches, pairwise comparisons of the fungal communities revealed differences between *Rinse-dip* and *Shake-cut* (Adonis test, Table S3b).

Alpha diversity indices, namely Observed and Chao, computed on bacteria ASV counts, did not show differences between the unplanted soil and the sampling approaches *Rinse-dip* and *Shake-dip* (Fig. S2a, c). Conversely, lower numbers of taxa were observed for the *Shake-vortex* and *Shake-cut,* indicating a more marked rhizosphere effect (ANOVA and Tukey post hoc test, Observed P-value = 1.59×10^-15^, Chao P-value = 2.67×10^-14^) (Fig. S2a, c). Fungal community analyses of the same indexes showed significantly greater richness in bulk unplanted soil compared to all sampling approaches (ANOVA and Tukey post hoc test, Observed P-value = 3.30×10^-5^, Chao P-value = 4.03×10^-5^) (Fig. S2b, d). *Shake-vortex* had the lowest Chao richness; however, this was significant just in comparison to *Shake-dip* (P-value = 0.04) (Fig. S2d). All sampling approaches had similar Observed fungal richness (P-value > 0.05) (Fig. S2b).

Considering the bacterial and fungal community diversity Shannon index, which accounts for the number of taxa and their abundances, all the rhizosphere sampling approaches showed a significant microbial diversity reduction compared to the unplanted bulk soil (ANOVA and Tukey post hoc test, bacterial 16S rRNA P-value = 1.28×10^-15^; Fungal ITS P-value =7.95×10^-10^) (Fig. 3d, e). For bacterial communities, excluding bulk unplanted soil, the Shannon index was greater in the *Rinse-dip* and *Shake-dip* approaches than in *Shake-vortex* and *Shake-cut*, with a significant increase observed for *Rinse-dip*.

We included a ‘dip-supernatant control’ to capture bacteria that may have remained in the supernatant due to insufficient sedimentation for the approaches *Rinse-dip* and *Shake-dip*. (the solution volume was decanted before centrifuging the remaining 50 mL). The ‘dip-supernatant control’ richness was significantly lower than in rhizosphere samples; however, it still contained >1000 taxa representing a consistent proportion (between 63 −75 %) of the number of taxa retrieved in the rhizosphere samples (Fig. S3).

Finally, we were interested in understanding whether any of the sampling approaches inadvertently introduced plant-derived 16S rRNA contamination into the rhizosphere samples. We compared the number of reads assigned as ‘Chloroplast’ as a proxy for plant contamination for each of the unplanted bulk soils and sampling approaches (Fig. S4, ANOVA and Tukey post hoc test, P-value = 5.29 x 10^-9^). The bulk unplanted soils samples displayed lower content of chloroplast reads than any of the rhizosphere samples (Fig. S4). Among the sampling approaches samples, the greatest number of reads assigned as ‘Chloroplasts’ was retrieved from *Rinse-dip*, whereas the lowest was assigned to *Shake-dip*. The number of chloroplast reads was not significantly different for *Shake-vortex* or *Shake-cut* compared to *Rinse-dip* or *Shake-dip* (Fig. S4).

### Different sampling approaches capture a conserved microbiota reflecting most of the microbial abundance

We performed pairwise comparisons to assess microbial abundances across different sampling approaches (Fig. 4, Table S4). In bacteria at ASV level, ‘vortexing’ approaches showed a remarkable percentage of rhizosphere reads enriched (38.14-25.32%), reflecting greater abundances of the enriched taxa compared with the ‘dipping’ approaches (15.86-1.34%) (Table S4a). These differences were reflected at the phylum level and clustered by ‘dipping’ and ‘vortexing’ approaches (Fig. 4a). The phyla enriched in ‘dipping’ approaches were less abundant than the phyla enriched in ‘vortexing’ approaches (Fig. 4a, b). Although differences in bacterial abundance were detected in pairwise comparisons, the majority of rhizosphere-enriched ASVs (349 ASVs) were recovered across all four sampling approaches, with a cumulative abundance of 77.67% rhizosphere reads and recapitulates the phyla composition of the rhizosphere with Proteobacteria, followed by Actinobacteria, Bacteroidota and Firmicutes (Fig. 4 b, Wald test Individual P-values < 0.05, FDR corrected).

**Fig. 4:**
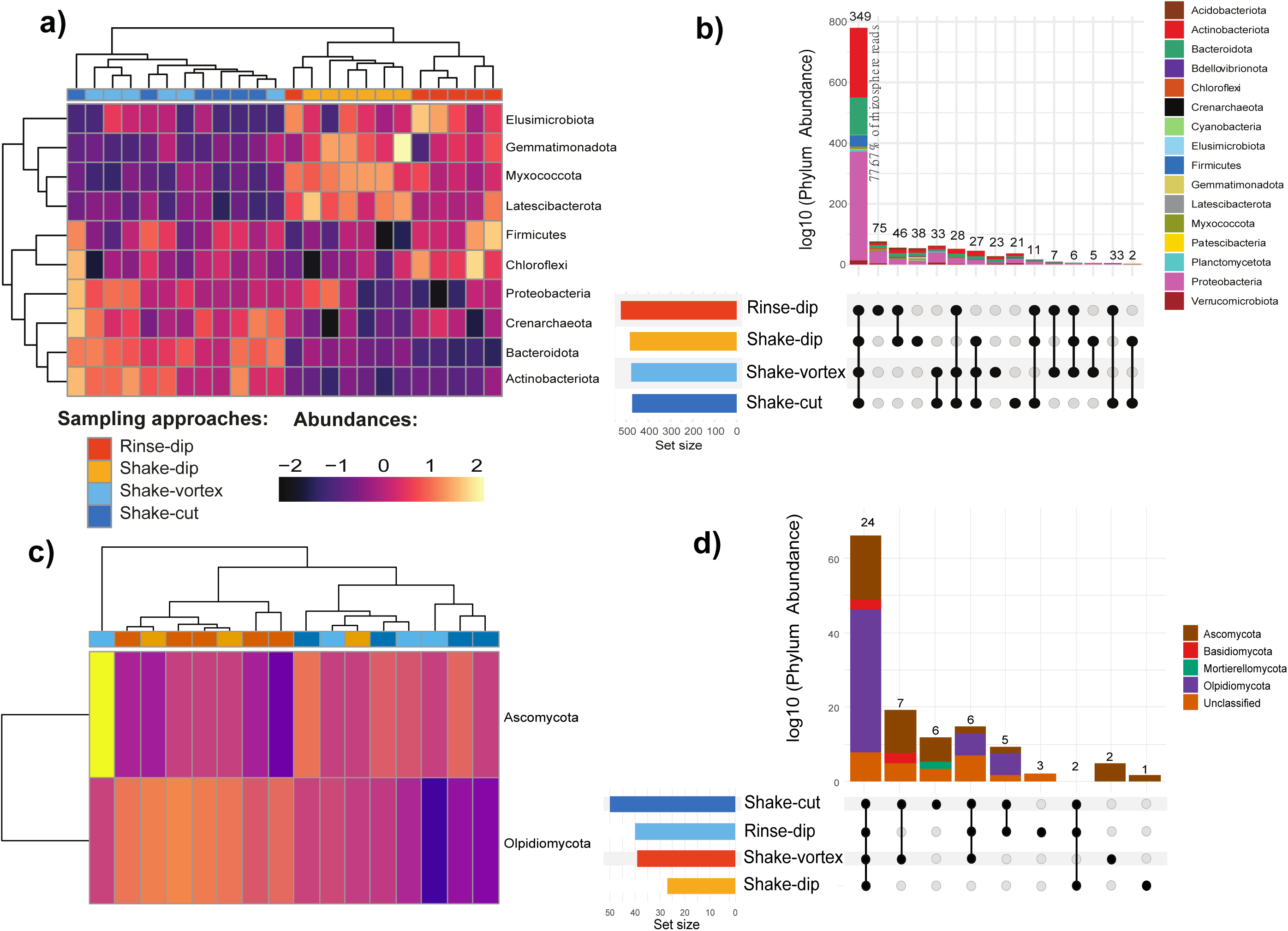
Heatmap of differential microbial enrichment at phylum level across protocols in (a) the bacteria 16S rRNA gene and (c) the fungal ITS. Phyla differentially enriched at individual p-values < 0.05, Wald Test, FDR corrected. UpSet plots of taxa (ASVs) simultaneously enriched in pairwise comparisons retrieved from the different rhizosphere sampling approaches in (b) the bacteria 16S rRNA gene and (d) the fungal ITS. Vertical bars denote the number of ASVs enriched shared or unique for each comparison depicted as relative abundance at phylum level, while the horizontal bars refer to the number of ASVs enriched in the indicated rhizosphere sampling approach. ASVs differentially enriched at individual p-values < 0.05, Wald Test, FDR corrected.

In fungi, the percentage of rhizosphere reads enriched at ASV level between sampling approaches (0-2.8%), indicates even less marked impact in the choice of sampling approach compared with bacteria (Table S4 b). Hierarchical clustering of the fungal taxa enriched from the rhizosphere revealed the presence of three main clusters (Fig. 4c). Two of the clusters differentiated *Rinse-dip* and *Shake-vortex*, while the third cluster contained one *Shake-cut* sample. *Shake-dip* and *Shake-cut* samples showed distinct profiles and were not clustering (Fig. 4c). The abundance of ITS fungal ASVs, like bacteria, (25 ASVs), shared 72.87 % of rhizosphere reads between approaches. Its phylum composition was dominated by Olpidiomycota, followed by a group of unclassified taxa, Ascomycota, and Basidiomycota (Fig. 4d Wald test individual p-values < 0.05, FDR corrected).

### The absolute abundances of microbial nitrogen cycle genes were unaffected by the sampling approach

Next, we investigated how sampling approaches impacted on functional groups of the microbiome. The abundance of the *amoA* gene (nitrification) and the *nosZII* clade II gene (denitrification) were not significantly affected by the sampling approach (Fig. S5). However, differences were identified between the unplanted soil and some of the sampling approaches (Fig. S5). Overall, we found a (mostly non-significant) trend of reduced nitrification-denitrification genetic potential in the rhizosphere compared to bulk soil. The *amoA* gene was significantly less abundant in *Shake-dip* compared with its cognate unplanted control *Bulk-dip*, whereas *nosZII* abundance was reduced in the rhizospheres of the sampling approaches *Shake-vortex* and *Shake-cut* compared with its unplanted control *Bulk-vortex* (Fig. S5).

## Discussion

In this study, we tested and evaluated different sampling approaches for root exudation and microbiota profiling from the same soil-grown plant. We observed differences in root biomass and morphology and, to a lesser extent, in the rhizosphere microbiota composition, among the sampling approaches tested. While we found minor differences in organic N root exudation, total dissolved C root exudation rates and non-targeted metabolite profiles were unaffected by the different sampling approaches. Our results benchmark a novel methodology for sampling root exudates and the rhizosphere microbiota, which enables increased sampling throughput.

### Sampling approach selection influences root biomass and morphology, and to a lesser extent, root exudation

After sampling, aboveground plant biomass was unaffected. However, below-ground biomass varied among the different sampling approaches, with *Rinse-dip* resulting in a greater root biomass than *Shake-vortex*. (Fig. 2b). As the mode of bulk soil removal (i.e. *Rinsing* vs *Shaking*) did not result in significant differences in root biomass, this indicates that ‘vortexing’ is more destructive than ‘dipping,’ inducing root losses. Interrelated root morphology traits, including root length, surface area, average diameter, and root area, followed the same trend, confirming that root biomass was lost during ‘vortexing’ (Table S1). ‘Vortexing’ has been widely used in sampling rhizosphere microbiota to detach the rhizosphere soil from the roots (Maver *et al*., 2021; Escudero-Martinez *et al*., 2022; Terrazas *et al*., 2022). However, if the goal is to assess root morphology, in which preserving root system integrity is essential, then ‘vortexing’ is not suitable for sampling. ‘Vortexing’ is also operationally limited to smaller and/or younger root systems. Yet, ‘vortexing’ also has important advantages such as faster sample throughput in comparison to the ‘dipping’ approach, and a smaller volume of solution containing the rhizosphere soil slurry needs to be processed. We did not find differences between bulk soil removal by ‘dipping’ or ‘shaking,’ but it is important to note that we used one soil type with a sandy texture in our study (Sandy Silt Loam). Different soil textures might have an impact on sampling (Barillot *et al*., 2013). Bulk soil removal via ‘shaking’ might cause more root loss and damage in heavier soils with larger clay contents due to break-off of large root-containing heavy soil aggregates in comparison to ‘rinsing.’

We used chloroplast reads in the rhizosphere as a proxy for plant content in the microbiota samples. The number of reads was similar across sampling protocols; only *Shake-dip* rhizosphere samples contained fewer chloroplast reads (Fig. S4). Therefore, even if ‘vortexing’ was associated with root biomass loss, this was insufficient to cross-contaminate the microbiota sample preparation.

Overall, the different sampling approaches showed a minor effect on barley exudation patterns. N root exudation rates were affected, while C exudation rates and the non-targeted metabolite composition were comparable (Fig. 2c-e). A recent study demonstrated that experimental factors such as the RSVR can bias the exudation results (Otxandorena-Ieregi *et al*., 2024). Here, we reported a lower RSVR in *Shake-vortex* due to root biomass loss, which may explain the differences observed in the N root exudation (Fig. S1). Although, Otxandorena-Ieregi et al. (2024) did not investigate N root exudation in their study, our results suggest that N root exudation rates might be more sensitive to the concentration gradient introduced by RSVR than C root exudation and non-targeted metabolomics (Fig. 2c, d).

Alternatively, the larger degree of root damage/root loss due to ‘vortexing’ might also have caused more release of N-containing metabolites. An early study already demonstrated that root cell damage and osmotic adjustment reactions can result in artificially enhanced root exudation rates (Ayers & Thornton, 1968). Root damage is unavoidable in the soil-hydroponic hybrid setup, irrespective of the sampling approach, but the degree of root damage can differ depending on the sampling approach. In our experimental set up, we included a short root adjustment step (see Materials and Methods: *<u>1.3. Exudate sampling)</u>* before placing the root system into the final sampling solution that should have captured the root cell damage while providing root osmotic adjustment (Otxandorena-Ieregi *et al*., 2024). The same authors reported that C root exudation rates remained comparable with an RSVR between 2 and 4 g root dry weight L^-1,^ which is in line with our experimental conditions and results. Nevertheless, our results indicate that, despite minor differences in N exudation, the exudation process is unaffected by the tested root system manipulations prior to the hydroponic exudate collection.

### Microbiota diversity showed a distinctive ‘rhizosphere effect’ in comparison to the unplanted soil, but it varies within sampling approaches

Sampling approaches did not affect the absolute abundances in the bacterial community but did impact the fungal abundances (*Shake-dip* had a lower abundance than *Shake-vortex*) (Fig. 3 b). This may be due to a better retrieval of fungal biomass by ‘vortexing’ the entire root system. However, these differences did not impact on fungal beta or alpha diversity.

All sampling approaches showed clear rhizosphere pattern across kingdoms (fungi and bacteria) in either alpha or beta diversity compared to the unplanted soil controls (Fig. 3). These observations confirm that the plant’s signature on microbiota assembly is sufficiently robust to withstand biases induced by different sampling approaches. However, there were differences among the rhizosphere soil sampling approaches (Fig. 3c, 3d). Microbiota composition was influenced by both the method used to remove bulk soil (rinse or shake) and the approach taken to harvest rhizosphere soil from roots (dipping or vortex), with more pronounced differences in bacterial communities than in fungal ones. Bacterial rhizosphere communities typically exhibit greater diversity and finer spatial heterogeneity than fungi (Schutes et al., 2025), meaning that small variations in soil handling, rhizosphere harvesting, or DNA extraction can disproportionately affect observed community composition.

An additional aspect that distinguishes each of these sampling approaches is how, once we harvested the rhizosphere soil from roots, the rhizosphere was collected: in ‘Vortexing’ sampling approaches (*Rinse-dip* and *Shake-dip*) the entire solution volume was centrifuged to retrieve the rhizosphere fraction from the solution, whilst in the ‘Dipping’ approaches (*Shake-vortex* and *Shake-cut*) we used gravitational sedimentation with the majority of the dipping solution being decanted after sedimentation and only the final 50 mL being centrifuged. Therefore, observed differences in results could be confounded by downstream differences in sampling steps. Perhaps centrifuging the entire dipping solution volume may be more critical for bacteria than for the fungal community, as marked differences were found between ‘Vortexing’ and ‘Dipping’ in the composition of this microbial community (Fig. 3 c). Indeed, we investigated the bacterial composition of the ‘dip-supernatant control’ for the ‘dipping’ sampling approaches (gravitational sedimentation). This control revealed that a fraction of the bacterial community was still present in the supernatant (Fig. S2). We did not attempt to amplify the supernatant of the ‘Vortexing’ sampling approaches (centrifugation). Taken together, this suggests that centrifuging samples is critical to harvest the microbial rhizosphere fraction, and it is a common step in rhizosphere sampling (Barillot *et al.,* 2013, Tkacz *et al.,* 2015). Conversely, alternative methods, such as gravitational sedimentation, may not recover the full microbial community, impacting both the microbial composition and abundance. This should be considered when designing subsequent experiments beyond the scope of this manuscript.

### A prominent ‘conserved barley microbiota’ was observed across the rhizosphere fractions for all sampling protocols

Due to the observed differences in microbial composition across sampling approaches (Table S3), we measured their impact on microbiota abundance for taxa enriched in the rhizosphere fraction driving the detected differences. Across the sampling protocols tested, both the bacteria and the fungal communities displayed a ‘core microbiota’ representing the greatest proportion of both, the numbers of taxa (16S: 349 ASVs and ITS: 25 ASVs) and the rhizosphere reads (16S: 77.67 % and ITS: 72.87 %) across sampling approaches (Fig. 4 b). The taxonomic composition and relative abundances of the sampling approaches’ bacterial core microbiome align well with previously reported phyla dominating the rhizosphere (Ling *et al*., 2022) and, in particular, the barley rhizosphere, such as Proteobacteria, Actinobacteria, and Bacteroidota (Escudero-Martinez and Bulgarelli, 2023). These bacterial phyla dominating the rhizosphere were even more abundant in ‘vortexing’ approaches (Fig. 4 a). Regardless the sampling approach used, the rhizosphere core fungal community, typically dominated by Ascomycota and Basidiomycota (Hennecke *et al.,* 2023), exhibits the largest relative abundance of the phylum Olpidiomycota. Although the dominance of this phylum in the rhizosphere of additional crops has been previously reported (Wei et al., 2023).

Altogether, this suggests that, regardless of the sampling protocol we proposed, a characteristic the rhizosphere microbial community profile was present in the enriched rhizosphere fractions. Therefore, despite variations in microbial composition across sampling approaches, rhizosphere microbiota sampling appeared more consistent and less variable than other plant-associated traits, such as root biomass and root morphology.

In conclusion, our results demonstrate that, under the tested conditions, different sampling approaches produced comparable microbiota and exudation patterns, enabling the integrated study of root exudation and microbial profiles from the same plant. We presented a detailed investigation of the implications of using different approaches to sample root exudates, which require minimal plant disturbance, and the rhizosphere microbiota fraction, which must be distinct from the unplanted soil. While the overall results were comparable, performing different sampling steps, i.e., rinsing, shaking, dipping, vortexing, gravitational sedimentation or centrifugation, introduced minor (but still statistically significant) differences in the obtained results. It is important defining the scope of the sampling and selecting the ‘sampling steps’ that will then best accomplish the experimental aim. Learning from the sampling approaches comparison presented here, we compiled a sampling method for subsequent experimentation that can be visualised in the format of a practice abstract (Additional information) and the following video: https://www.youtube.com/watch?v=Rkif6BbJmGc

## Supporting information

Primers, cycling conditions, and qPCR standard curves

---

**Fig. S1:**
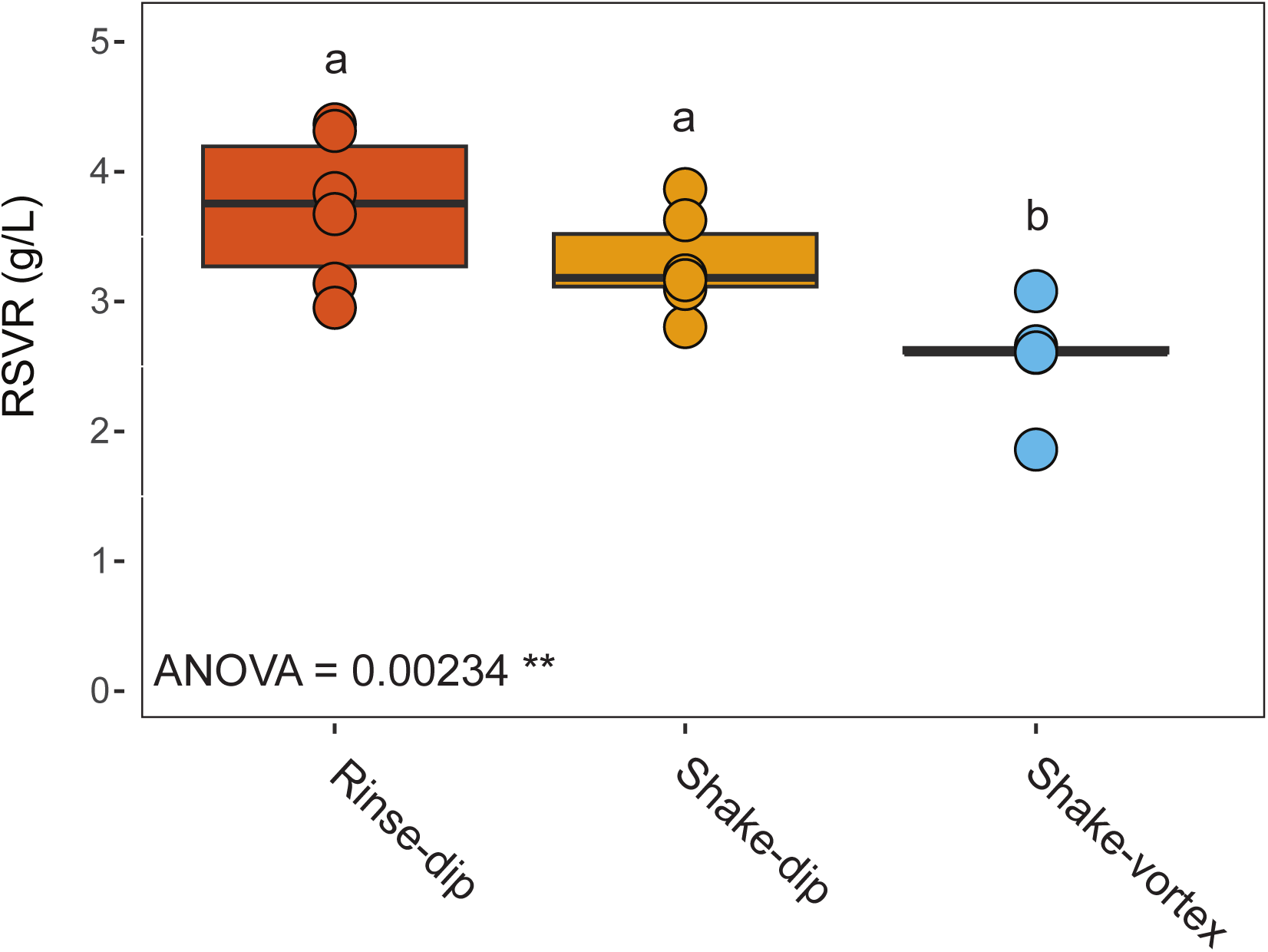
Box plots depicting the root (dry) weight-to-sampling solution volume (L) ratio (RSVR) obtained for the different sampling approaches. Individual dots depict individual biological replicates. Upper and lower edges of the box plots represent the upper and lower quartiles, respectively. The bold line within the box denotes the median. Letters indicate significant differences following ANOVA and post-hoc Tukey test; p-value<0.05.

**Fig. S2:**
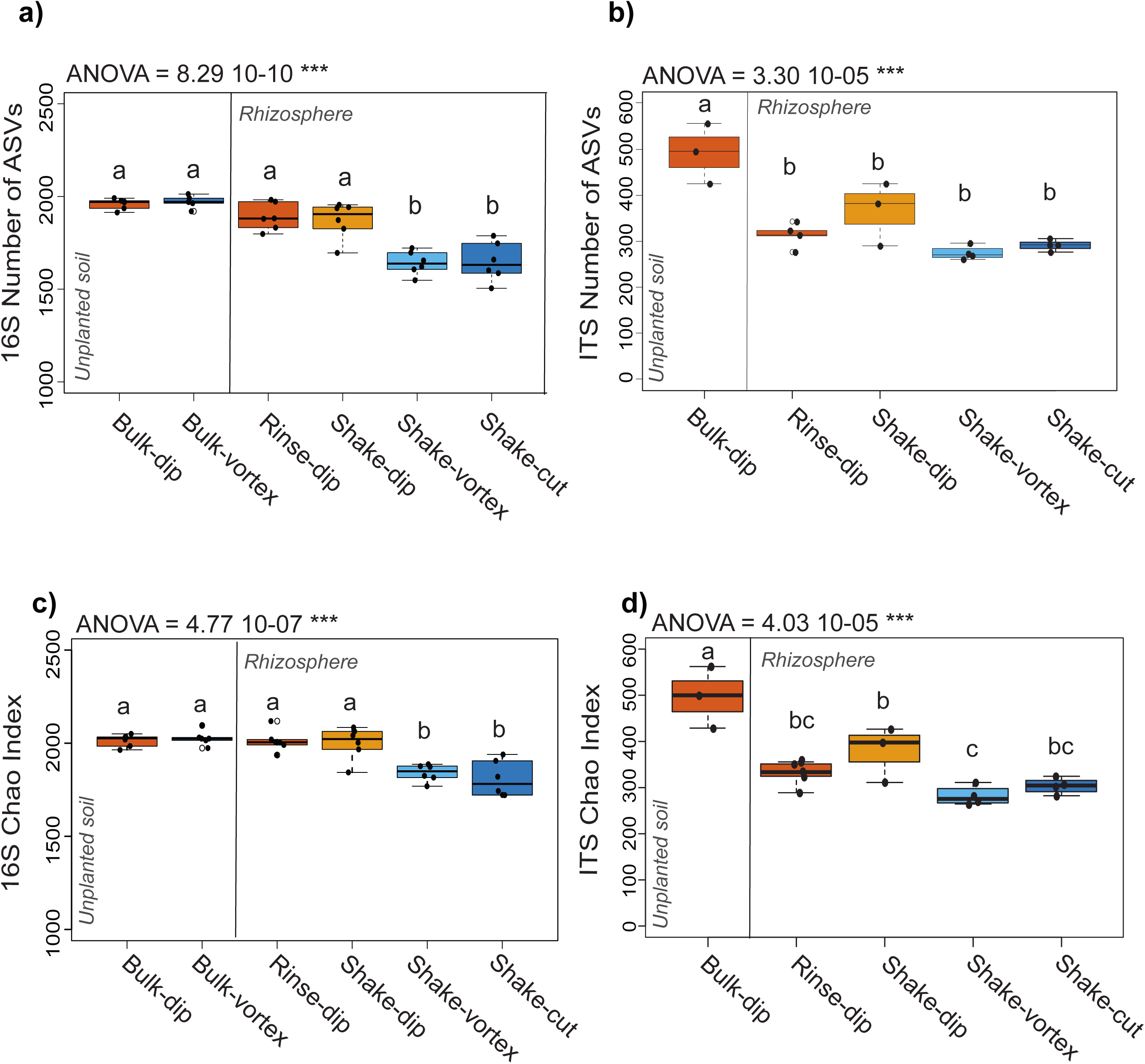
Boxplots representing the alpha diversity community richness (Observed and Chao) of the sampling approaches and the unplanted soil. Observed richness across sampling approaches and unplanted soil in bacteria (a) and fungi (c). Chao richness across sampling approaches and unplanted soil in bacteria (b) and fungi (d). In each panel, individual dots depict individual biological replicates. Upper and lower edges of the box plots represent the upper and lower quartiles, respectively. The bold line within the box denotes the median. Orange bulk soil samples (*Bulk-dip*) are the controls for *Rinse-dip* and *Shake-dip*, while blue bulk soil (*Bulk-vortex*) samples are the controls for *Shake-vortex* and *Shake-cut*. Letters indicate significant differences following ANOVA and post-hoc Tukey test; p-value<0.05.

**Fig. S3:**
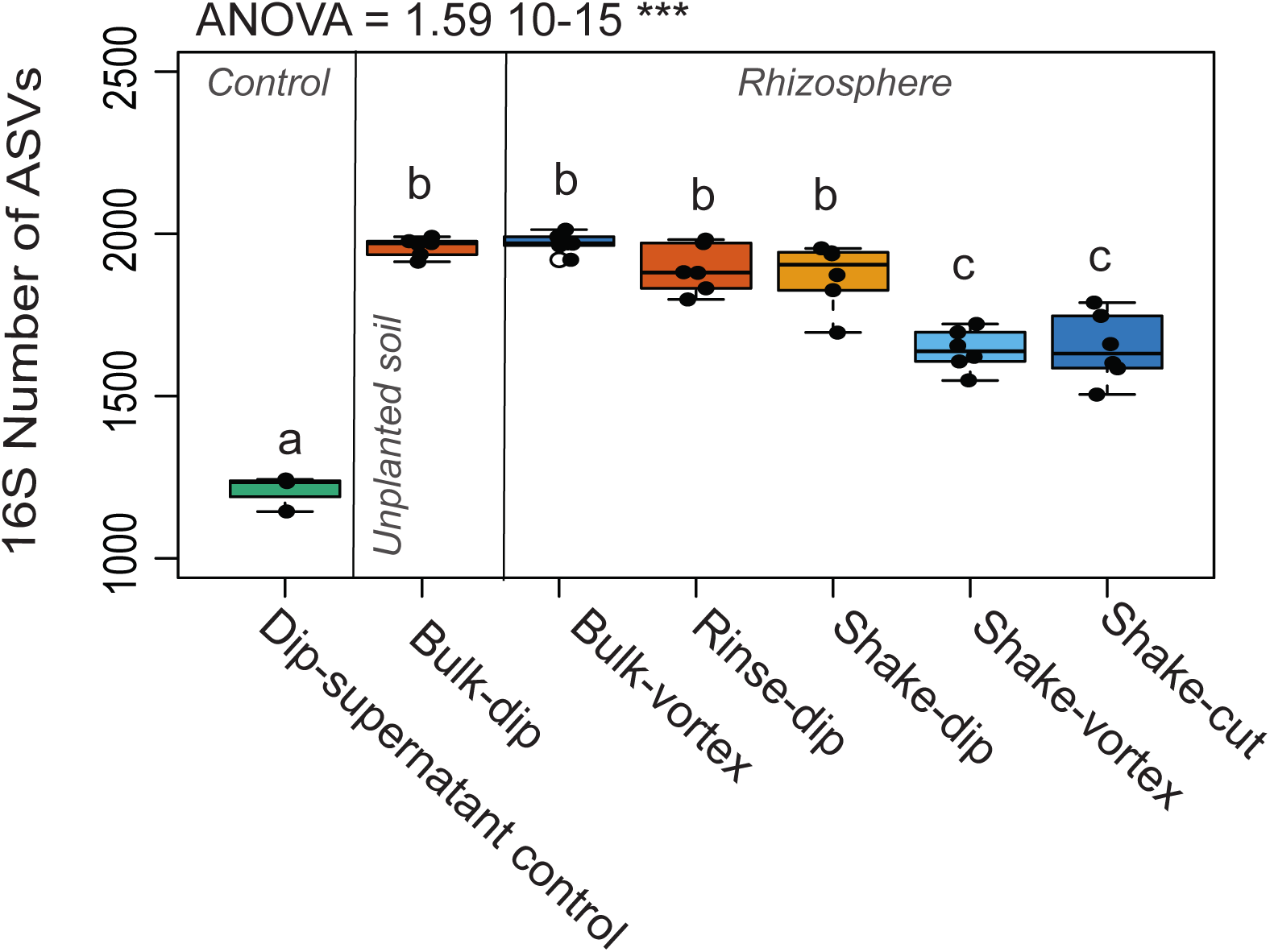
Boxplots representing the observed community richness of the supernatant control (green), the unplanted soil controls, and sampling approaches. In each panel, individual dots depict individual biological replicates. Empty dots are outliers. Upper and lower edges of the box plots represent the upper and lower quartiles, respectively. The bold line within the box denotes the median. Orange bulk soil samples (*Bulk-dip*) are the controls for *Rinse-dip* and *Shake-dip*, while blue bulk soil (*Bulk-vortex*) samples are the controls for *Shake-vortex* and *Shake-cut*. Letters indicate significant differences following ANOVA and post-hoc Tukey test; p-value<0.05.

**Fig. S4:**
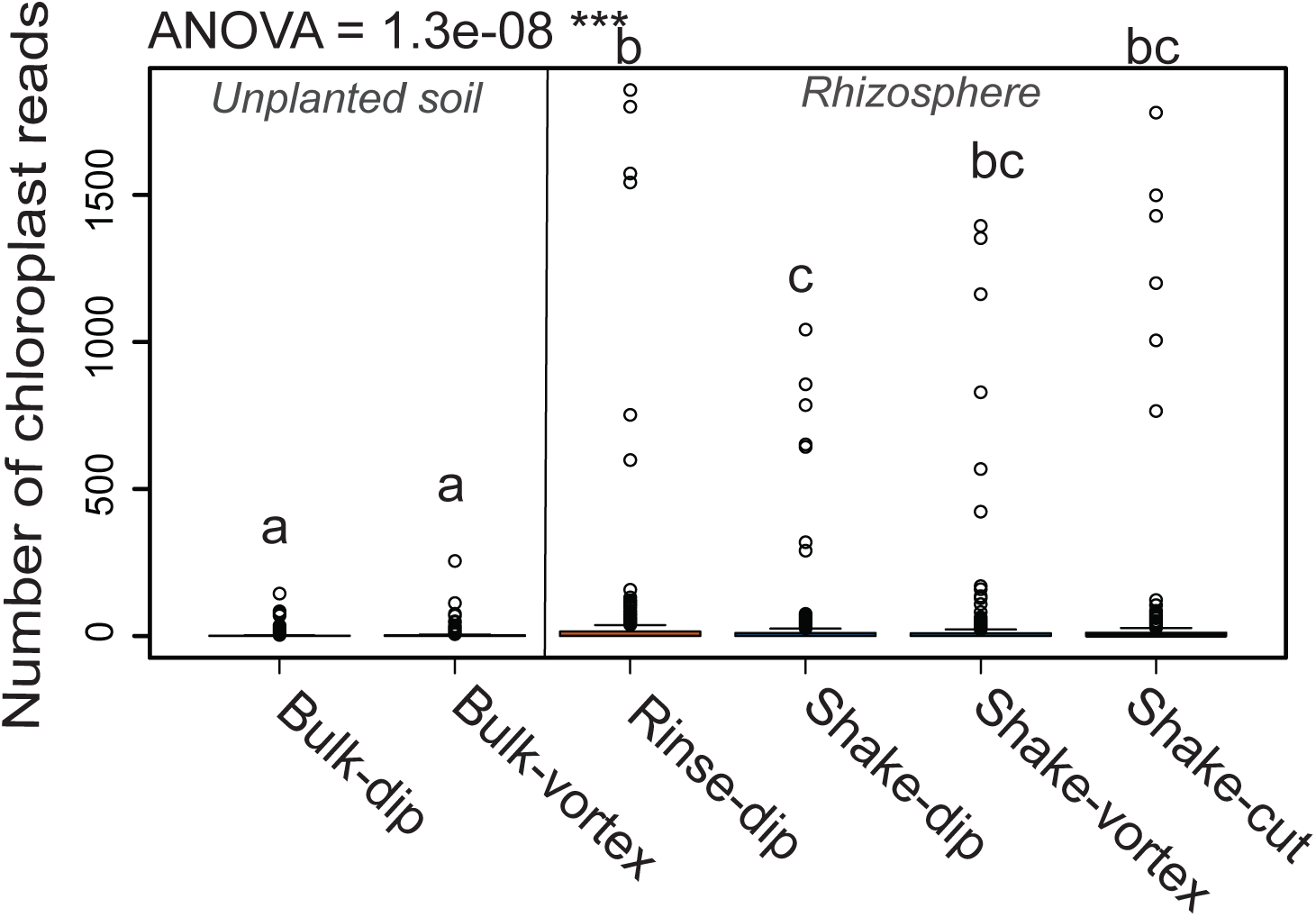
Box plot showing the number of reads assigned to chloroplast as a proxy for plant tissue contamination across the different sample types of rhizosphere fractionation methods and the bulk unplanted soil. Individual points represent the number of taxa classified as chloroplast (n=58) across samples. *Bulk-dip* soil samples are the controls for *Rinse-dip* and *Shake-dip*, while *Bulk-vortex* soil samples are the controls for *Shake-vortex* and *Shake-cut*. Sample differences were assessed with ANOVA and post-hoc Tukey test; p-value<0.05.

**Fig. S5:**
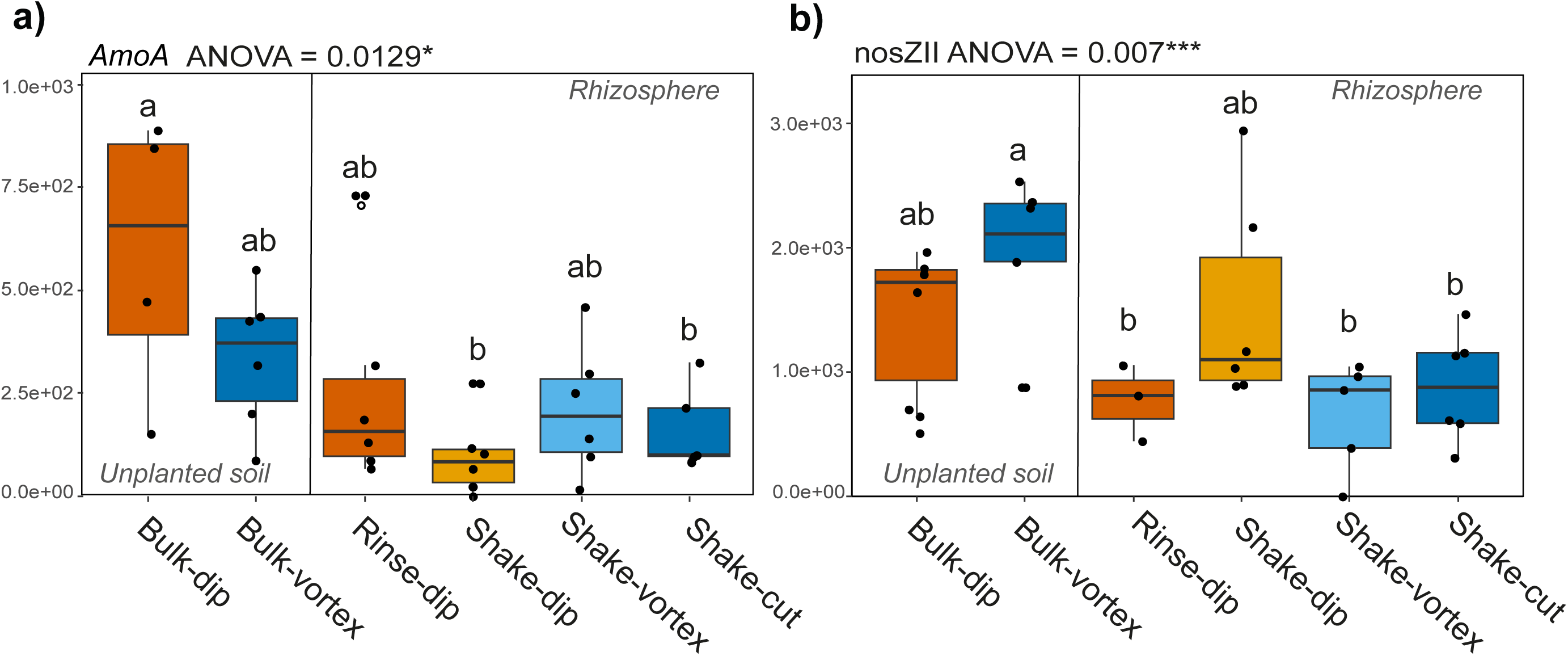
Microbiota quantification of nitrogen cycling microbial genes. (a) *AmoA* gene (nitrification) and (b) *noz2* clade II (denitrification) gene copies per ng of DNA in unplanted soil and rhizospheres obtained with different sampling approaches. Box plots depict the total copy numbers of each nitrogen cycling gene per ng of DNA. Orange bulk soil samples (Bulk-dip) are the controls for Rinse-dip and Shake-dip, while blue bulk soil (Bulk-vortex) samples are the controls for Shake-vortex and Shake-cut. Individual dots depict individual biological replicates. Different letters denote significantly different groups (ANOVA and post hoc Tukey HSD test).

**Table S1:** Root morphology parameters recorded after handling the plant samples with different sampling approaches. The values represent the average values and the standard error of the mean (mean±SE). Differences were assessed using an ANOVA and post-hoc Tukey test. Letters denote significant differences.

| **Protocol** | **Root Length (m)** | **Root Surface Area (cm<sup>2</sup>)** | **Average Diameter (mm)** | **Specific Root Length** | **Projected Area (cm<sup>2</sup>)** |
| --- | --- | --- | --- | --- | --- |
| ***Rinse-dip*** | 3,702 ±358.4 **a** | 434 ±42.8 **a** | 1.12 ±0.1 **a** | 12,440 ±7.8 **a** | 138.03±13.6 **a** |
| ***Shake-dip*** | 3,214 ±201.2 **ab** | 356 ±21.6 **ab** | 1.03 ±0.5 **ab** | 12,181 ±4.2 **a** | 113.37±6.9 **ab** |
| ***Shake-vortex*** | 2,432±129.2 **b** | 267 ±16.0 **b** | 1.01 ±0.1 **b** | 12,032 ±8.9 **a** | 84.90±5.1 **b** |

**Table S2:**
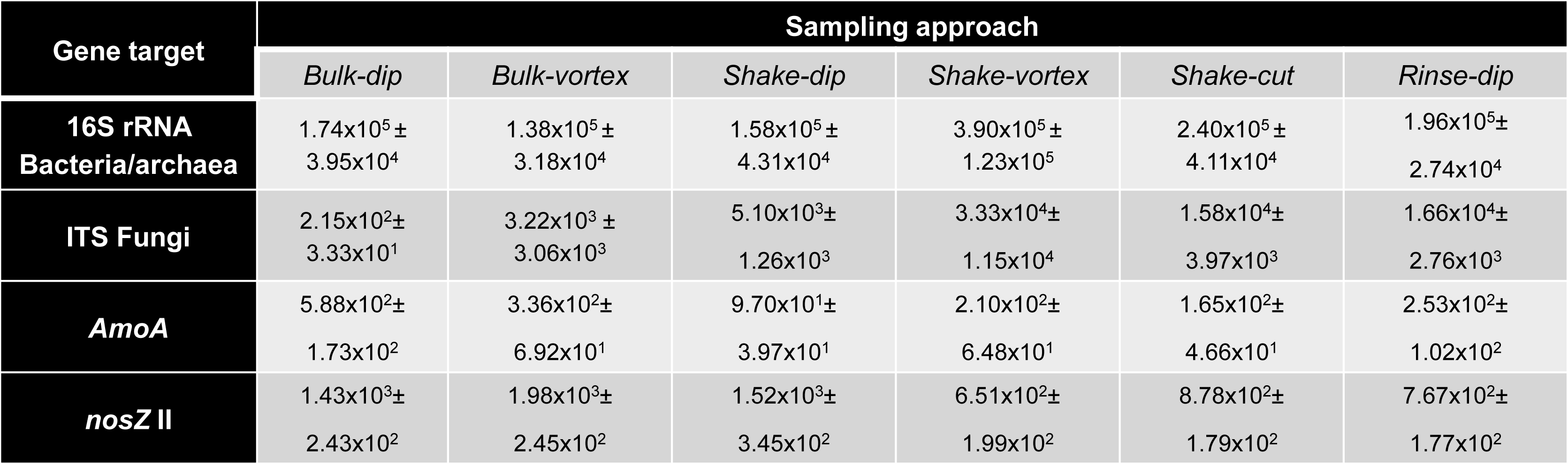
Results of qPCR determination of gene abundances of selected taxonomic and nitrogen cycling genes. The values represent the average values and the standard error of the mean (mean±SE).

| Gene target | Sampling approach |  |  |  |  |  |
| --- | --- | --- | --- | --- | --- | --- |
|  | *Bulk-dip* | *Bulk-vortex* | *Shake-dip* | *Shake-vortex* | *Shake-cut* | *Rinse-dip* |
| **16S rRNA**<br>**Bacteria/archaea** | 1.74x10<sup>5</sup>±<br>3.95x10<sup>4</sup> | 1.38x10<sup>5</sup>±<br>3.18x10<sup>4</sup> | 1.58x10<sup>5</sup>±<br>4.31x10<sup>4</sup> | 3.90x10<sup>5</sup>±<br>1.23x10<sup>5</sup> | 2.40x10<sup>5</sup>±<br>4.11x10<sup>4</sup> | 1.96x10<sup>5</sup>±<br>2.74x10<sup>4</sup> |
| **ITS Fungi** | 2.15x10<sup>2</sup>±<br>3.33x10<sup>1</sup> | 3.22x10<sup>3</sup> ±<br>3.06x10<sup>3</sup> | 5.10x10<sup>3</sup>±<br>1.26x10<sup>3</sup> | 3.33x10<sup>4</sup>±<br>1.15x10<sup>4</sup> | 1.58x10<sup>4</sup>±<br>3.97x10<sup>3</sup> | 1.66x10<sup>4</sup>±<br>2.76x10<sup>3</sup> |
| ***AmoA*** | 5.88x10<sup>2</sup>±<br>1.73x10<sup>2</sup> | 3.36x10<sup>2</sup>±<br>6.92x10<sup>1</sup> | 9.70x10<sup>1</sup>±<br>3.97x10<sup>1</sup> | 2.10x10<sup>2</sup>±<br>6.48x10<sup>1</sup> | 1.65x10<sup>2</sup>±<br>4.66x10<sup>1</sup> | 2.53x10<sup>2</sup>±<br>1.02x10<sup>2</sup> |
| ***nosZ II*** | 1.43x10<sup>3</sup>±<br>2.43x10<sup>2</sup> | 1.98x10<sup>3</sup>±<br>2.45x10<sup>2</sup> | 1.52x10<sup>3</sup>±<br>3.45x10<sup>2</sup> | 6.51x10<sup>2</sup>±<br>1.99x10<sup>2</sup> | 8.78x10<sup>2</sup>±<br>1.79x10<sup>2</sup> | 7.67x10<sup>2</sup>±<br>1.77x10<sup>2</sup> |

**Table S3:** Table illustrating the percentage of variance explained in pairwise comparisons between sampling approaches in a) bacteria and b) fungi. The number in the table shows the proportion of variance (*R*^2^) explained by the factor ‘Sampling approach’ for rhizosphere samples. The asterisks associated with the *P-value* denote its significance by the Adonis test.

|  |  |  |  |  |
| --- | --- | --- | --- | --- |
| **a)** | **R<sup>2</sup>-P-value** | ***Shake-dip*** | ***Shake-vortex*** | ***Shake-cut*** |
|  | ***Rinse-dip*** | R2 = 0.233,<br>P-value = 0.012* | R2 = 0.450,<br>P-value = 0.018* | R2 = 0.394,<br>P-value = 0.012* |
|  | ***Shake-dip*** | - | R2 = 0.427,<br>P-value = 0.036* | R2 = 0.373,<br>P-value = 0.006** |
|  | ***Shake-vortex*** | - | - | R2 = 0.108,<br>P-value = 1 |
| **b)** | **R<sup>2</sup>-p-value** | ***Shake-dip*** | ***Shake-vortex*** | ***Shake-cut*** |
|  | ***Rinse-dip*** | R2 = 0.217,<br>P-value = 1 | R2 = 0.689,<br>P-value = 0.084 | R2 = 0.663,<br>P-value = 0.03* |
|  | ***Shake-dip*** | - | R2 = 0.278,<br>P-value = 0.150 | R2 = 0.252,<br>P-value = 0.396 |
|  | ***Shake-vortex*** | - | - | R2 = 0.105,<br>P-value = 1 |

**Table S4:** Microbial differential abundance between protocols expressed in taxa (ASV) (upper table) numbers and percentage of reads (lower table) for the a) bacteria 16S and b) the fungal ITS. ASVs differentially enriched at individual p-values < 0.05, Wald Test, FDR corrected.

| Number of 16S rRNA ASVs | *Rinse-dip* | *Shake-dip* | *Shake-vortex* | *Shake-cut* |
| --- | --- | --- | --- | --- |
| *Rinse-dip* | - | 67 | 9 | 83 |
| *Shake-dip* | 15 | - | 15 | 22 |
| *Shake-vortex* | 123 | 146 | - | 0 |
| *Shake-cut* | 101 | 164 | 2 | - |

| % reads 16S rRNA | *Rinse-dip* | *Shake-dip* | *Shake-vortex* | *Shake-cut* |
| --- | --- | --- | --- | --- |
| *Rinse-dip* | - | 7.41 % | 1.34 % | 1.72 % |
| *Shake-dip* | 2.43 % | - | 13.35 % | 15.86 % |
| *Shake-vortex* | 38.14 % | 25.32 % | - | 0 % |
| *Shake-cut* | 33.63 % | 27.86 % | 0.33 % | - |

b)
| Number of ITS ASVs | *Rinse-dip* | *Shake-dip* | *Shake-vortex* | *Shake-cut* |
| --- | --- | --- | --- | --- |
| *Rinse-dip* | - | 0 | 8 | 4 |
| *Shake-dip* | 3 | - | 7 | 5 |
| *Shake-vortex* | 5 | 2 | - | 4 |
| *Shake-cut* | 2 | 0 | 1 | - |

| % reads ITS | *Rinse-dip* | *Shake-dip* | *Shake-vortex* | *Shake-cut* |
| --- | --- | --- | --- | --- |
| *Rinse-dip* | - | 0% | 0.2% | 0.2% |
| *Shake-dip* | 0.3% | - | 0.9% | 0.8% |
| *Shake-vortex* | 0.002% | 0% | - | 0% |
| *Shake-cut* | 0% | 2.8% | 0% | - |

**Supplementary Information 1:** Primers, standard curves, and cycling conditions used in this investigation.

## Acknowledgements

This work was supported by the following research grants: Horizon Europe: Root2Resilience (https://root2res.eu, grant agreement N°101060124); ERC Starting Grant Phyto Trace (project number 801954), DFG, German Research Foundation (project number 403803214); Austrian Science Fund (project number FWF I 4445); OTR15203 Project Talent Attraction from Salamanca Ciudad de Cultura y Saberes foundation and Salamanca Council, CLU-2025-2-02 Unit of Excellence IRNASA_CSIC, from Junta Castilla y León and EU FEDER and Project DEEP-MaX-2024_IRNASA funded by CSIC. In addition, the Scottish Government Research Programme supports the work of the James Hutton Institute staff.

## Author contributions

DB, TSG and FB and EO funding acquisition, project administration, mentoring, resources, and supervision. CEM and EO conceptualization, methodology and supervision. CEM, EB, HS, MS, MB, LB, DR, JM, and EO investigation. CEM, EB, HS, MS, DA, JM, PH, PT, JA and EO data curation and formal analysis. CEM, EB and EO writing-original draft. All authors writing, review and editing.

## Data availability

The Sequences generated in this study will be made public upon acceptance for publication of the preprint in the European Nucleotide Archive (ENA), accession number PRJEB97221. Scripts to reproduce figure and statistical analyses are available at https://github.com/carmenmariaescudero and

